# Quantitative differences between intra-host HCV populations from persons with recently established and persistent infections

**DOI:** 10.1101/2020.06.17.157792

**Authors:** Pelin Icer Baykal, James Lara, Alex Zelikovsky, Yury Khudyakov, Pavel Skums

## Abstract

**Background:** Detection of incident hepatitis C virus (HCV) infections is crucial for identification of outbreaks and development of public health interventions. However, there is no single diagnostic assay for distinguishing recent and persistent HCV infections. HCV exists in each infected host as a heterogeneous population of genomic variants, whose evolutionary dynamics remain incompletely understood. Genetic analysis of such viral populations can be applied to the detection of incident HCV infections and used to understand intra-host viral evolution.

**Methods:** We studied intra-host HCV populations sampled using next-generation sequencing from 98 recently and 256 persistently infected individuals. Genetic structure of the populations was evaluated using 245,878 viral sequences from these individuals and a set of selected parameters measuring their diversity, topological structure, complexity, strength of selection, epistasis, evolutionary dynamics, and physico-chemical properties.

**Findings:** Distributions of the viral population parameters differ significantly between recent and persistent infections. A general increase in viral genetic diversity from recent to persistent infections is frequently accompanied by decline in genomic complexity and increase in structuredness of the HCV population, likely reflecting a high level of intra-host adaptation at later stages of infection. Using these findings, we developed a Machine Learning classifier for the infection staging, which yielded a detection accuracy of 95.22%, thus providing a higher accuracy than other genomic-based models.

**Interpretation:** The detection of a strong association between several HCV genetic factors and stages of infection suggests that intra-host HCV population develops in a complex but regular and predictable manner in the course of infection. The proposed models may serve as a foundation of cyber-molecular assays for staging infection, that could potentially complement and/or substitute standard laboratory assays.

**Funding:** AZ and PS were supported by NIH grant 1R01EB025022. PIB was supported by GSU MBD fellowship.

## 1. Introduction

Hepatitis C virus (HCV) infection remains a major cause of morbidity and mortality, with an estimated 70 million people being HCV infected worldwide in 2015^1^. HCV infection is the leading cause of chronic liver diseases and hepatocellular carcinoma worldwide, contributing to the death of more than 350,000 people in 2015^1^ outbreaks continue to occur, posing a serious challenge to public health^2^. HCV is highly mutable. As a result, each infected individual hosts a heterogeneous population of genetically related HCV variants or *quasispecies^3^*. Substantial diversity of intra-host viral populations plays a crucial role in disease progression and epidemic spread^4–6^. However, intra-host dynamics of HCV and other RNA viruses remain poorly understood. One of the most important questions is the relative contribution of random and deterministic evolutionary factors in disease progression, or, using the metaphor of S.J. Gould^7^, whether is it possible to “replay the tape of life” for the virus evolution inside a host. This question is of high importance for biomedical research, as predictability of viral evolution potentially implies the power to understand and control the disease^8,9^, which may result in advanced diagnostic and treatment strategies.

In this paper, we study evolutionary factors associated with the transition between HCV infection stages. In more than 50% of cases untreated HCV infection proceeds to the chronic phase, which can lead to the development of liver cirrhosis and/or hepatocellular carcinoma^8^. Accurate recent or persistent staging of HCV infection is important for biomedical applications. In clinical settings, it may inform the patient management and treatment strategy. In epidemiology, identification of acute cases allows for detection and investigation of recent transmissions and outbreaks and provides information on disease incidence. Understanding of changes in intra-host HCV populations at different stages of infection would constitute a large step towards reliable forecasting of viral evolutionary dynamics.

Recent HCV infection is usually accessed using clinical symptoms and time since seroconversion. HCV infection may, however, remain asymptomatic for years while seroconversion is not frequently detected, preventing accurate identification of infection stages. Several laboratory methods have been reported for distinguishing acute and chronic stages of infection^10,11^. Detection of HCV RNA in the absence of anti-HCV activity in serum specimens was used as an indication of recent HCV infection^12^. Although a strong marker, it has a very short duration and cannot be used for reliable detection of acute infections.

Advent of next-generation sequencing (NGS) presented an opportunity to sample and analyze unprecedented large numbers of intra-host viral variants from numerous infected individuals. HCV variants sampled by NGS have been used to detect stages of HCV infection^13,14^. The stage detection methods are generally based on the assumption that intra-host viral evolution is driven by the continuous immune escape resulting in genetic diversification. Consequently, quantitative measures of genetic diversity of intra-host viral variants are assumed to be most useful for staging. However, several recent reports contested the veracity of this assumption. In particular, after initial diversification, intra-host HCV populations may actually lose heterogeneity and stop diverting at later stages of infection,^5,15^ with certain viral variants persisting in infected hosts for years^5,16^. Furthermore, this process is accompanied by increase of negative selection over the course of HCV infection^5,15,17,18^. These findings suggest a high level of intra-host adaptation at late stages of infection^4^ and in.dicate that genetic heterogeneity is not a reliable marker for infection staging, and more elaborate metrics are needed to understand HCV evolution and to accurately classify recent and persistent HCV infection.

Here, we present a new approach for staging HCV infection using quantitative genomic measures to evaluate diversity, information content, effective dimensionality, topological structure, evolutionary dynamics and physical-chemical properties of intra-host HCV variants and populations. Analysis of parameters’ distributions at early and late stages of infection suggests that intra-host HCV populations evolve in a complex but regular and predictable manner. Based on these findings, we propose a multi-parameter machine learning classifier for staging HCV infection. The model allows for more accurate detection of recent HCV infection than models based only on population diversity and provides new insights into mechanisms of infection progression.

## 2. Materials and Methods

### 2.1 Data Collection and Preprocessing

We analyzed intra-host HCV populations sampled from recently (N=98) and persistently (N=256) infected persons collected as described in^35^. The E1/E2 junction of the HCV genome (*L* = 246nt), which contains the hypervariable region 1 (HVR1), was sequenced using the GS FLX System and the GS FLX Titanium Sequencing Kit (454 Life Sciences, Roche, Branford, CT). Obtained sequences were processed using the error correction and haplotyping algorithm KEC^19^, which produced 245,878 unique viral haplotypes with frequencies.

### 2.2 Parameters Calculation

The analyzed parameters could be loosely split into four groups: genomic parameters, complexity parameters, network parameters and biochemical parameters. We assumed that a given intra-host population contains *n* unique haplotypes with frequencies *f*_1_, …, *f*_n_. Sixteen parameters corresponding to this population constitutes its *feature vector*.

### Genomic Parameters

These parameters are obtained by direct comparison of sequences from each population. *Distance-based parameters* include *mean and standard deviation* of pairwise hamming distance distribution, and the *conservation score* of the population consensus sequence calculated with the NUC44 scoring matrix. We also utilized the so-called *mutation frequency* parameter,^13^ which is defined as the mean distance between all haplotypes and the most frequent haplotype. All four parameters measure the population diversity.

Diversity was also quantified using three *entropy-based parameters*. For a genomic position *i*, its positional *k*-entropy is defined as the entropy of the frequency distribution of *k*-mers (subsequences of length *k*) starting at *i*. An *average positional k-mer entropy E_k_* is the mean of positional *k*-entropies over all positions:

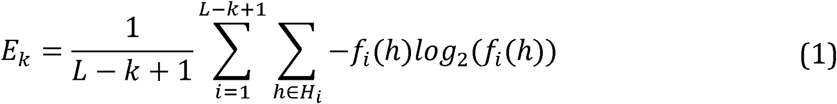

Here *h* is a *k*-mer, *H*_i_ is the set of *k*-mers starting from the *i*-th position and *f*_*i*_(*h*) refers to the relative frequency of h inside *H_i_*. For *k* = *L* the parameter *E*_*L*_ is an entropy of observed haplotype frequencies, while for *K* = 1 it is an average position-wise nucleotide entropy. In our model we used entropies *E*_1_, *E*_*L*_ and *E*_10_.

Next, we estimated the frequency of *transversions* (mutations between purines and pyrimidines) among all observed mutations within the population. This parameter is suggested by previous studies^20^ that reported higher frequencies of transitions over transversions in HCV populations. *Selective Pressure* was measured using the DN/DS ratio, which has been calculated as the ratio of rates of non-synonymous (DN) and synonymous (DS) substitutions with respect to the most frequent genomic variant.

### 2.3 Complexity Parameters

*PCA complexity* is derived from principal component analysis (PCA). For each population, its alignment is transformed into *n* × *L* numerical matrix, and the complexity is defined as the percentage of principal components required to explain at least α = 50% percent of the observed genetic variance. PCA complexity measures the effective dimensionality of the population as the multidimensional system.

*Kolmogorov complexity* is the classical concept of information theory, which quantifies the descriptive/information complexity of a string over a finite alphabet. Informally it is defined as the highest possible degree of compression of a given string without loss of information. Although the exact value of Kolmogorov complexity is algorithmically incomputable, it can be efficiently approximated using data compression techniques. In our case, each viral sequence has been transformed into a binary string, the strings have been concatenated, and Kolmogorov complexity of the resulting string has been estimated by a variant of Lempel-Ziv algorithm^21^.

### 2.3.1 Network Parameters

This group of parameters is derived from the analysis of *genetic networks* of HCV populations, that represent a *sequence space^20^* of a virus. Formally, for each patient its genetic network *G_N_ = (V, E)* is a graph, whose vertices *V* represent sampled viral haplotypes, and edges E connect variants which differ by at most *T* mutations (by default *T* = 1) (see Fig. 1). With each vertex we associate the frequency of the corresponding haplotype. In the case of a large population size accompanied by a high mutation rate and a fast reproduction time, genetic networks constructed using NGS data represent population structures significantly more accurately than phylogenetic trees^17^. Their structure is shaped by various factors, such as epistasis, founder effects, and selection pressures that affect the virus over the course of infection^5,22^. For each network, the following four parameters have been calculated.

**Figure 1:**
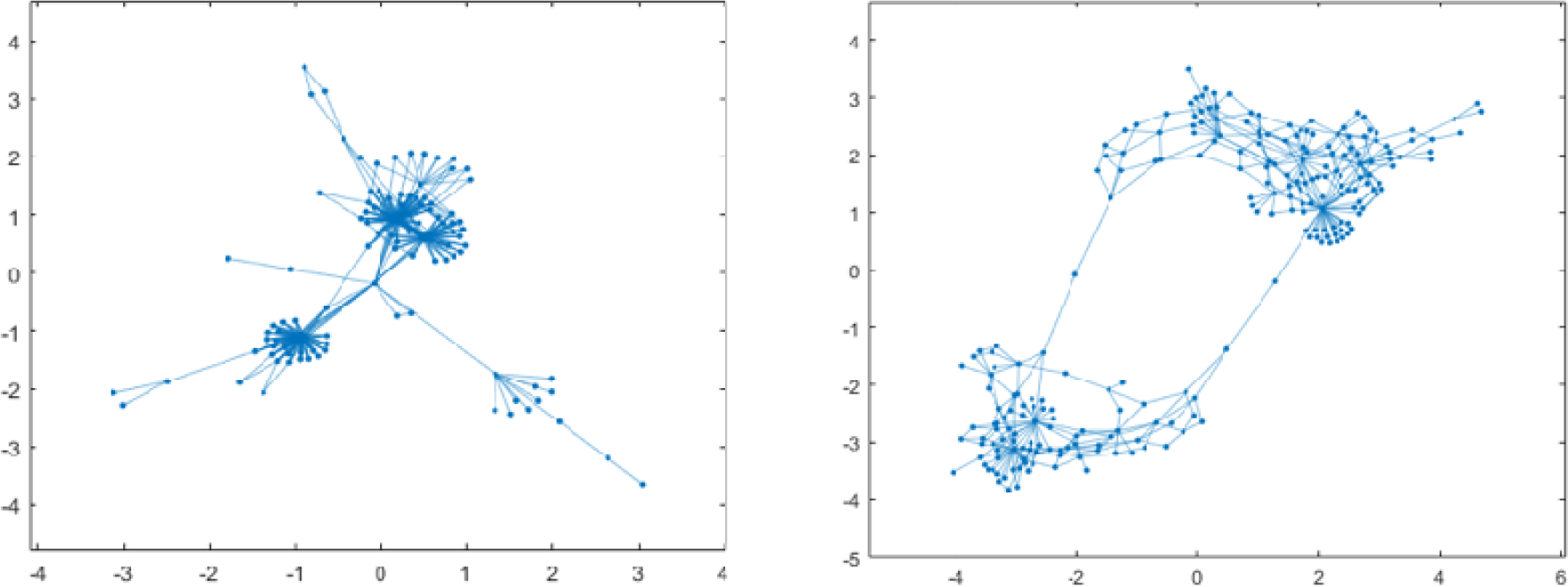
Examples of genetic viral networks for a recently infected (left) and a persistently infected (right) individual. The viral network of the recently infected host has the structural properties typical for scale-free networks.

*Robustness/selection balance* has been measured by the correlation between vectors of vertex frequencies and eigenvector centralities. The latter is the principal eigenvector of the adjacency matrix of *G_N_*. In the classical quasispecies model, vertex centralities are indicative of the mutational robustness of corresponding viral variants^23^, while a high frequency may be indicative of a higher fitness.

Topological structures of genetic networks have been assessed using two parameters. The first of them is *s-metric^24^* 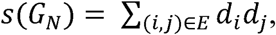, which measures how close a network is to being scale-free (here, *d*_*i*_ is a degree (number of neighbors) of a vertex _*i*_). Scale-free networks are ubiquitous in biological and social systems and share specific properties such as a power-law degree distribution, small diameter and presence of hubs. To account for variable sample sizes, is normalized by the factor (the order of magnitude of the maximum-metric for-vertex network).

The second network structural parameter is the clustering coefficient, which measures the degree to which network vertices tend to cluster together. It is defined as the probability that a random connected vertex triplet is complete (i.e. every pair of vertices is connected by an edge).

*Evolutionary dynamics parameter* estimates an age of the genetic network using an evolutionary model. Given *n* viral variants, we simulate their frequencies using a system of ordinary differential equations (S1)–(S3), which describes the interaction of the viral population with the host’s immune system (Supplemental Section S1). We classify populations as recent or persistent based on the qualitative behavior of the function describing the deviation of simulated and observed frequencies over time. Formally, we define an estimated population age as the time, when simulated viral frequencies achieve the best agreement with observed frequencies, i.e., where is a Jensen-Shannon divergence between distributions and. Owing to the inherent uncertainty of the quantitative parameters of the model, rather than using as a prediction variable we utilize qualitative characteristics of the divergence function. Namely, persistent and recent populations are characterized by divergence functions with descending and ascending trends, respectively (Fig. S1, see Supplemental Section S1). The classification is performed separately for each connected component of the genetic network, and the patient is classified as persistently infected (the parameter), if at least one of the components is persistent, and as recently infected (, otherwise).

### 2.3.2 Biochemical parameter

For each viral sequence, we assess whether this sequence has physico-chemical properties associated with recent or persistent infection. The biochemical index of an intra-host population is thus defined as the sum of frequencies of variants identified as having a physico-chemical profile pointing to persistent infection.

The method for evaluation of the properties of a given viral haplotype is described in detail in Supplemental Section S2. Briefly, for a given sequence we construct its biochemical feature profile using the following physico-chemical indexes of DNA dimers: the thermodynamic indexes (Breslauer-dH and Breslauer-dG), structural indexes (twist-tilt, slide-rise, protein-DNA twist, slide-2, twist-1), the nucleotide composition index (G-content) and the energy indexes of DNA (stabilizing energy of Z DNA and enthalpy)^25^. Such set of parameters can accurately measure changes in structure-function relationships and can be used to predict a broad range of biological and biochemical properties of DNA/RNA biomolecules^25^. The constructed set of features is processed by the problem-specific dimensionality reduction and feature selection pipeline, and binarized. The obtained binary feature vector representations of intra-host HCV variants were used as input data to train a stochastic gradient descent (SGD) classifier^26^. The SGD classifier implements regularized linear models with stochastic gradient descent (SGD) learning and is a very efficient approach, with linear training cost, which can easily be scaled to big data problems. Selection and tuning of the hyperparameters of the SGD classifier was done using a balanced training set (1,968 and 1,965 feature vectors for sequences sampled from recently and persistently infected hosts) and assessed by five-fold cross-validation.

### 2.4 Machine Learning Classifier

Feature vectors of recently and chronically infected hosts were used to train machine learning classifiers for infection stage prediction. Given a labeled training set comprising feature vectors together with their class labels (recent or persistent), each classifier is fitted to the training data by adjusting its model parameters and assigns labels for unlabeled feature vectors using the trained model. In this study, we used Support Vector Machines with linear and polynomial kernel and Logistic Regression. Both approaches are classical supervised learning methods that construct a hyperplane in the multidimensional Euclidean space, which serves as a separator for feature vectors from classes of recently and persistently infected hosts.

## 3 Results

### 3.1 Stage-specific distributions of parameters

Except for several diversity measures (*k*-entropy, site entropy, mean distance, conservation score and mutation frequency), there is a small-to-medium correlation between the parameters (Fig. 2), demonstrating that they reflect different properties of intra-host viral populations.

**Figure 2:**
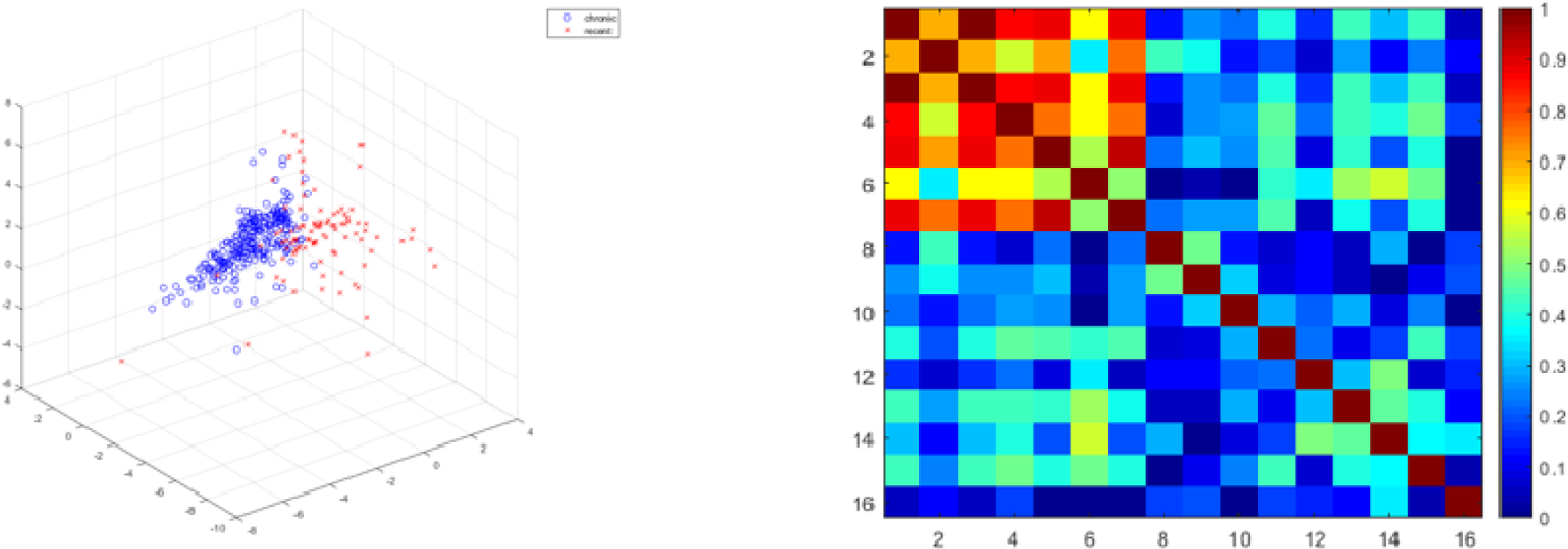
Left: 3-D projection of feature vectors of recently and persistently infected hosts (with highly correlated feature removed) constructed by multidimensional scaling. Right: heatmap of absolute values of pairwise correlations betwee parameters.

Feature vectors of recent and persistent populations are separable from each other (Fig. 2). For each parameter, Mann-Whitney U-test suggests statistically significant difference between recent and persistent intra-host populations (Table 1).

**Table 1:** Parameters with a significant association to the infection stage. The columns contain p-values of U-test, mean values, and 95% confidence intervals for viral populations among persistently and recently infected persons.

| Parameters | p-value | Persistently infected |  | Recently infected |  |
| --- | --- | --- | --- | --- | --- |
|  |  | Mean | 95% CI | Mean | 95% CI |
| 1. Mean distance | 1.41E-24 | 0.034 | (0.031, 0.037) | 0.015 | (0.012, 0.017) |
| 2. Std distance | 4.10E-14 | 0.019 | (0.018, 0.021) | 0.010 | (0.007, 0.013) |
| 3. Conservation score | 5.11E-25 | 0.422 | (0.393, 0.451) | 0.188 | (0.155, 0.221) |
| 4. Mutation frequency | 3.07E-16 | 0.023 | (0.021, 0.026) | 0.010 | (0.007, 0.013) |
| 5. $k$-entropy | 7.00E-23 | 0.630 | (0.602, 0.658) | 0.357 | (0.323, 0.392) |
| 6. Frequency entropy | 4.56E-06 | 0.668 | (0.648, 0.689) | 0.567 | (0.527, 0.607) |
| 7. SNV entropy | 1.14E-21 | 0.084 | (0.079, 0.089) | 0.043 | (0.037, 0.048) |
| 8. Transversion mutation | 1.06E-07 | 0.061 | (0.054, 0.068) | 0.032 | (0.027, 0.038) |
| 9. DN/DS | 5.39E-10 | 0.713 | (0.626, 0.779) | 1.330 | (1.096, 1.565) |
| 10. PCA complexity | 4.91E-04 | 0.014 | (0.012, 0.026) | 0.034 | (0.023, 0.045) |
| 11. Kolmogorov complexity | 1.55E-11 | 0.041 | (0.040, 0.043) | 0.052 | (0.048, 0.056) |
| 12. Robustness/Selection | 3.66E-15 | 0.628 | (0.608, 0.647) | 0.386 | (0.329, 0.442) |
| 13. $s$-metric | 1.93E-20 | 0.001 | (0, 0.002) | 0.044 | (0.023, 0.065) |
| 14. Clustering coefficient | 2.08E-13 | 0.082 | (0.064, 0.100) | 0.356 | (0.292, 0.420) |
| 15. ODE | 2.42E-06 | -0.270 | (-0.378, -0.162) | 0.224 | (0.055, 0.394) |
| 16. Biochemical parameter | 2.92E-35 | 0.628 | (0.614, 0.642) | 0.379 | (0.354, 0.403) |

As expected, diversities are on average higher for persistent than recent populations (*p*-values between 1.41 · 10^−24^ and 4.56 · 10^−6^; Fig. 3 (1-7)). Higher genetic diversity of persistent populations is accompanied by significantly lower PCA and Kolmogorov complexities (*p* = 4.91 · 10^−4^ and *p* = 1.55 · 10^−11^; Fig. 3 (10-11)). This could be explained by the role of intra-host adaptation during the later stage of infection, when genomes are highly specific to the environment and SNVs selected over the course of intra-host evolution are highly interdependent, thus reducing the effective dimensionality of the population. It is known that high Kolmogorov complexity indicates high level of randomness of a sequence, while low complexity implies the presence of specific structural patterns inside a sequence. Thus, lower Kolmogorov complexity at later stages of disease suggests the increase in strength of epistatic connectivity among nucleotide positions during intra-host evolution and points to a higher level of adaptation and specialization of members of intra-host populations. At the earlier stages of infection, nucleotide changes are seemingly more random, resulting in populations with higher dimensionality. Increase in negative selection additionally contributes to the reduction of dimensionality and complexity at later stages of HCV infection (*p* = 5.39 · 10^−10^; Fig. 3 (9)).

Transition mutations were overwhelmingly more frequent than transversion mutations for both classes of samples. This fact agrees with the previously published results^20^, although the magnitude of difference vary along the genome: HVR1 transitions are times more frequent than transversions, while a 75-fold difference was reported for NS5B^20^. Prevalence of transversions was times higher in persistent populations (*p* = 1.06 · 10^−7^; Fig. 3 (8)). This phenomenon could be interpreted as another reflection of increasing intra-host adaptation over the course of infection. Indeed, transversions represent a higher genetic barrier for the selection of escape mutants from HCV-specific immune responses^20^. Thus, growth of transversion frequencies at later evolutionary stages may mark a declining role of immune escape and a growing role of other evolutionary mechanisms such as adaptation by antigenic cooperation^4^.

**Figure 3:**
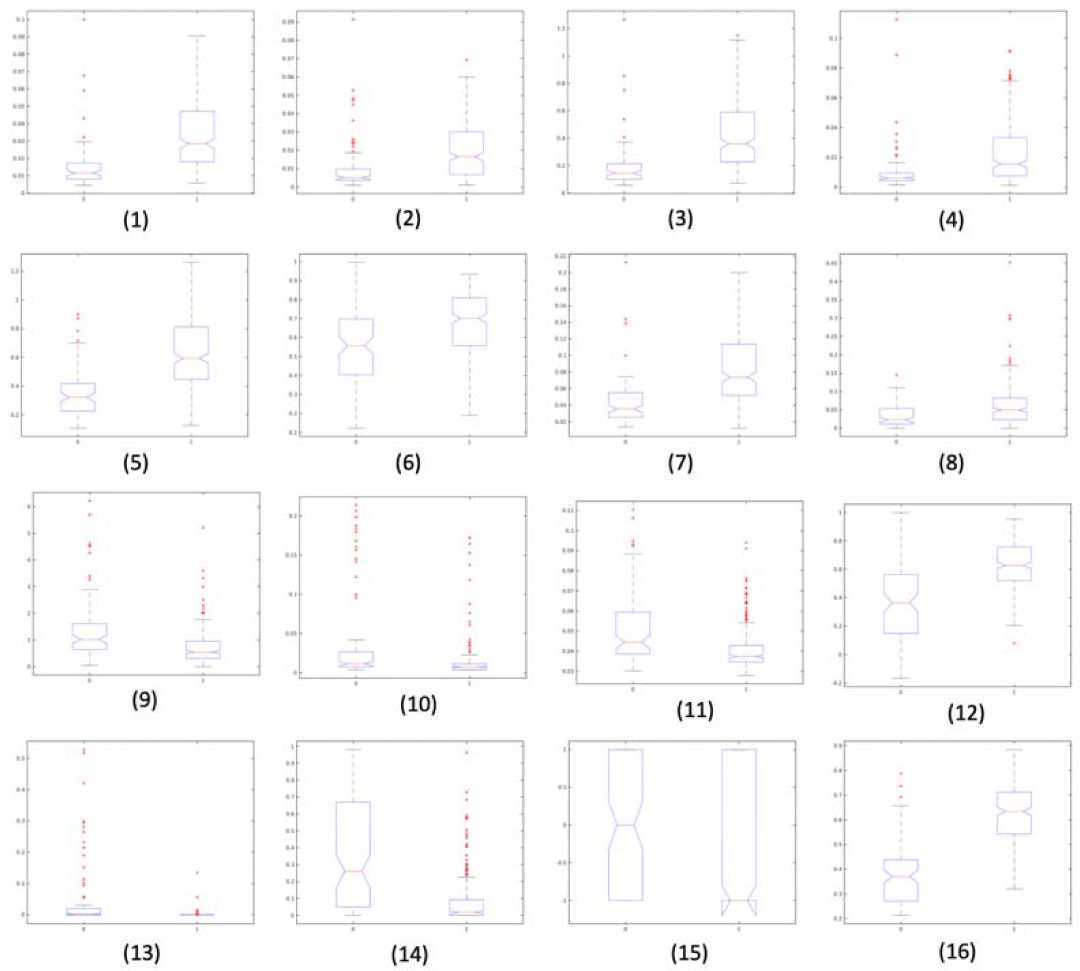
Box plots of parameter distributions for recent (left box plot on each graph) and persistent (right box plot on each graph) intra-host HCV populations. The plots are in the same order as in Table 1.

Genetic networks of recent and persistent intra-host populations possess different structural properties. Networks of recent populations have significantly higher *s*-metrics and clustering coefficients (*p* = 1.93 · 10^−20^ and 2.08 · 10^−13^; Fig. 3 (13-14)). It indicates that, in contrast to the persistent populations, they tend to have structural properties more typical for scale-free networks, including the power-law degree distribution with clearly manifested hubs (high-degree vertices), with their vertices having propensity to cluster (Fig.1). This observation can be explained by the role of founder viral variants at the earlier stage of infection. A significantly higher correlation between frequencies and network centralities of variants in persistent populations (*p* = 3.66 · 10^−15^; Fig. 3 (12)) indicates that the population structure at later stages is significantly influenced by mutational robustness, while at earlier stages it is basically defined by founders. Recent and persistent HCV populations are also separable by an evolutionary dynamic parameter *c_ODE_* (*p* = 2.42. 10^−6^; Fig. 3 (15)).

Finally, individual sequences of recent and persistent populations have distinct physico-chemical properties (*p* = 2.92 · 10^−35^; Fig. 3 (16)). It suggests that the physico-chemical property of HVR1 is influenced by, and is responsive to, within-host environmental factors specific to the recent and persistent stages of HCV infection.

### 3.2 Machine Learning Classification

Mutation frequency, *k*-entropy and frequency entropy have been excluded from the prediction model as they are highly correlated with other parameters. The remaining 13 parameters were used to train Support Vector Machines (SVM) and Logistic Regression classifiers for binary classification of intra-host viral populations labelled as “persistent” and “recent”. Accuracy of classifiers has been assessed using a two-step cross-validation. First, to account for the bias associated with unequal numbers of cases with persistent (*n* = 256) and recent (*n* = 98) infection, repeated random subsampling of 98 populations from the persistent sample dataset was performed. For each of the balanced training sets 10-fold cross-validation was carried out.

The average prediction accuracies are reported in Table 2. Classification performance evaluation of all methods indicates a high accuracy of infection stage inference, with SVM with quadratic kernel demonstrating the highest accuracy of 95.22%.

**Table 2:**
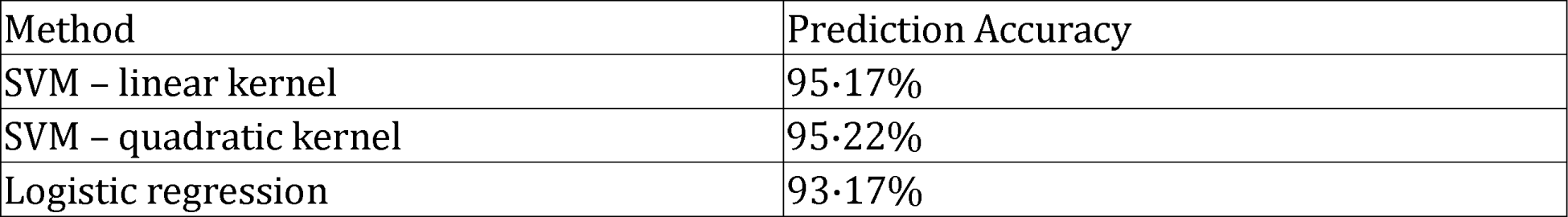
HCV infection stage prediction accuracies of machine learning methods

| Method | Prediction Accuracy |
| --- | --- |
| SVM – linear kernel | 95.17% |
| SVM – quadratic kernel | 95.22% |
| Logistic regression | 93.17% |

SVM classifier with quadratic kernel has been compared to the previously published HCV infection staging models^13^ which classify intra-host viral populations as recent or persistent using frequency entropy, SNV entropy or mutation frequency. The ROC curves of the classifiers are shown in Fig. 4. Previously proposed methods (*AUROC* = 0,81, 0.66 and 0.78, respectively) were less accurate in comparison with the SVM classifier (*AUROC* = 0.99), thus suggesting that diversity parameters alone are not sufficient for accurate distinction between recent and persistent cases. SVM classifier performed at the expected lower accuracy on randomly labelled datasets (average *AUROC* = 0.4966), thus indicating that the associations between parameter distributions and infection stages are likely due to the structural and evolutionary factors rather than to random statistical correlations in the data.

**Figure 4:**
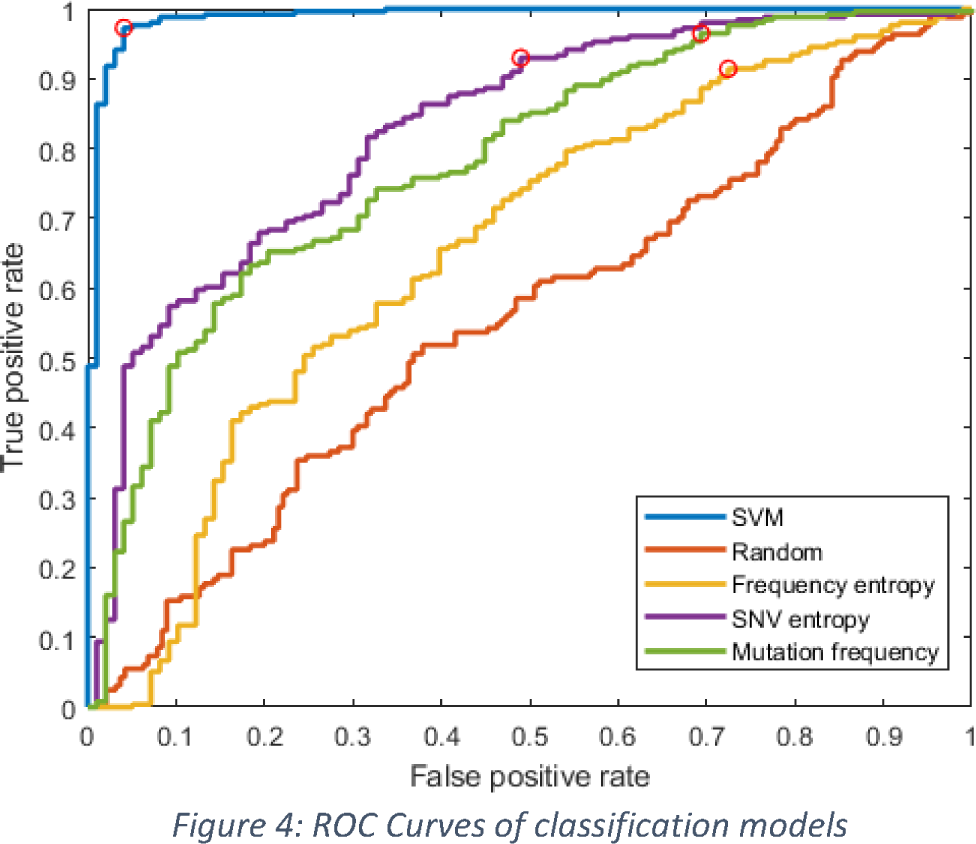
ROC Curves of classification models.

## 4 Discussion

We present the results of comprehensive analyses of the structure of intra-host viral populations using a large set of samples from individuals with recent and persistent infection, which significantly exceeds data sets used in earlier studies^13^. Amplicons covering HCV HVR1 have been sequenced by NGS. Intrinsically disordered regions (IDR) of proteins like HVR1 seem to be most useful for application in models to identify viral clinical properties from sequences. It has an extensive epistatic connectivity across the entire HCV polyprotein^27^, and is associated with immune escape^28^, drug resistance^27,29^ and virulence^30^. Consequently, IDRs play an important role in viral adaptation to the host environment, making regions like HVR1 sensitive “sensors” that accurately reflect intra-host biological changes during the infection process.

We identified a set of quantitative characteristics of intra-host HCV populations strongly associated with stages of infection. Our results indicate significant differences in the structure of HCV populations sampled from recently and persistently infected hosts. Models constructed using these parameters allowed us to train machine learning classifiers capable of inferring infection stage from HCV sequence data with accuracies as high as 95%. Our study confirms apreviously established positive correlation between infection stage and intra-host viral diversity^13,10,31^. However, because of complexities in the structural development of intra-host populations affected by bouts of selective sweeps and negative selection during chronic infection^4,32^, simple metrics of genetic heterogeneity are insufficient for the accurate staging of HCV infections. High accuracy could be achieved by using a combination of parameters measuring different structural and evolutionary properties of viral populations.

The proposed prediction models may serve as *cyber-molecular assays* for staging infection, that could potentially complement and substitute standard laboratory assays. In particular, the proposed models are currently being incorporated into Global Hepatitis Outbreak and Surveillance Technology (GHOST)^33^ — a web-based molecular surveillance system developed and maintained by CDC. They could also be applicable to other highly mutable viruses, such as HIV.

The detection of a strong association between several HCV genetic factors and stages of infection suggests that intra-host HCV populations develop in a complex but regular and predictable manner during the course of infection. Decline in dN/dS, increase in abundance of transversion mutations and decline in information complexity of HCV population progressing from the recent to persistent state is consistent with an orderly process of HCV population development during infection as was suggested earlier^4,5^ and is different from a model of an “arms race” predicting a continuous genetic diversification. These observations support the hypothesis that intra-host viral populations may evolve as quasi-social systems by complementary specialization of viral variants engaged in a certain type of cooperation^4,34^. Such specialization enables HCV populations to adapt to an intra-host environment as a group of cooperators rather than independent variants.

## Supporting information

Supplemental Section

## Ethics approval and consent to participate

Research was conducted as approved by the Institutional Review Board of the Centers for Disease Control and Prevention, Atlanta, GA (protocol 7270.0).

## Disclaimer

The findings and conclusions in this report are those of the authors and do not necessarily represent the official position of the Centers for Disease Control and Prevention and Georgia State University.

## Availability of data and material

The proposed method’s scripts are available in the following GitHub repository https://github.com/compbel/recentvschronic. The data can be requested from the CDC.

## Competing interests

The authors declare they don’t have any competing interests.

## Notes

### Competing Interest Statement

The authors have declared no competing interest.

