## Supplemental Section for "Quantitative differences between intra-host HCV populations from persons with recently established and persistent infections"

**S1. Model for calculation of evolutionary dynamics parameter.**

Viral population is described by the dynamical system (S1)-(S4), partially inspired by the ideas from^S1,S2^. Variables $c$, $c_{i}$, $v_{i}$, $r_{i}$, $h(i=1, \ldots, n)$ represent numbers of uninfected target cells, cells infected by virions with $i$th genome, virions with $i$th genome in the host's serum, $B$-cell antibodies targeting the *i*th genome and cytotoxic $T$-cells (CTLs), respectively. Target cells proliferate by a logistic growth law due to the limited liver cell carrying capacity. Cells are infected by a variant $i$ at rate $v_{i}$ and, after being infected, are eliminated by CTLs at rate $\delta h$. The genomes mutate at rate $\epsilon$, and the virions with the *i*th genome are introduced to the blood by the cells infected by the variant $j$ at the rate $pq_{ji}c_{j}$, where $q_{ji}={(\epsilon/3)}^{d_{ij}}{(1-\epsilon)}^{L-d_{ij}}$ is a probability of mutation between variants $i$ and $j$, whose genomes are at Hamming distance $d_{ij}.$ $B$-cell antibodies are variant-specific, while CTLs are cross-immunoreactive.   We used the values of model parameters estimated in^S1,S4^.

$$\dot{c}=\alpha+\rho\left( 1-\frac{c+\sum_{i=1}^{n} c_{i}}{C^{*}} \right)-\theta c-\beta c\sum_{i=1}^{n} v_{i}$$

(S1)

$$\dot{c}_{i}=\beta cv_{i}-\delta hc_{i}$$

(S2)

$$\dot{v}_{i}=p\sum_{ij\in E} q_{ji}c_{j}-\lambda r_{i}v_{i}$$

(S3)

$$\dot{r}_{i}=\phi v_{i}-\sigma r_{i}$$

(S4)

Simulated viral frequencies $g_{i}\left( t \right)$ are calculated as $g_{i}\left( t \right)=\frac{v_{i}(t)}{\sum_{i=1}^{n} v_{i}(t)}$ .

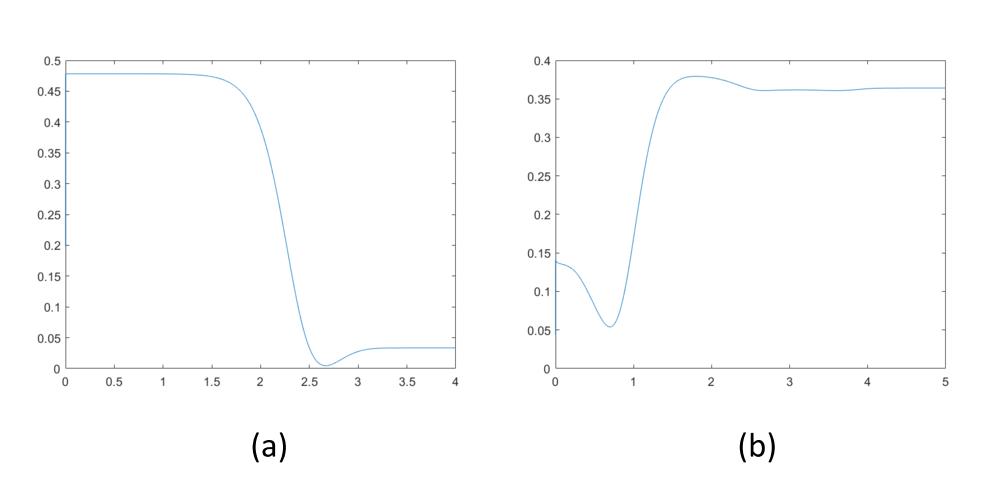

Figure S1: Example of function(t)for chronically infected ((a)) and recently infected ((b)) host

**S2. Calculation of biochemical parameter.**

To evaluate properties of a given sequence, we utilized the feature construction method^S4^ to represent the biochemical parameter space of nucleotide sequence variants of HCV HVR1. The features are based on following physico-chemical indexes of DNA dimers^S5^: the thermodynamic indexes (Breslauer-dH and Breslauer-dG), structural indexes (twist-tilt, slide-rise, protein-DNA twist, slide-2, twist-1), the nucleotide composition index (G-content) and the energy indexes of DNA (stabilizing energy of Z DNA and enthalpy)^25,S6^. Next, feature vector representations of HVR1 were constructed using dinucleotide-based auto-covariance formula $DAC'(u,Lag)$^S7^, where $u$ is a physico-chemical index and $Lag$ is the distance separating two nucleotide dimers along the nucleotide sequence. There are over 140 physico-chemical indexes with which to represent DNA dimers^S5^. To reduce the computation time and the size of feature vector representations (given by $u \times lag)$^S7^, we measured the Pearson correlation coefficient of each index $u$ to the chronic and recent classes to choose the best class-correlated indexes. Here, the physico-chemical index $u$ was represented by ten significantly correlated physico-chemical descriptors (R threshold value $\geq0.457$ and $p<2.20\cdot{10}^{-16}$). The auto-covariance per physico-chemical index $u$ was measured between nucleotide dimers separated by a distance of up to 60 nucleotides. To optimize the feature space representation of HVR1 in relation to the persistent/recent classes, we further processed the feature vectors by feature selection techniques^S8^. First, the class-attribute interdependence maximization (CAIM) algorithm^S9^ was used to automatically generate binary indicators for each of physico-chemical values in the feature vectors. Then, the CAIM-derived binary feature vectors were processed by the correlation-based feature selection (CFS) algorithm, which finds the most class-informative parameters based on a Merit scoring^S10^ (here, a feature subset of 54 variables with Merit score=0.6). CFS-based binary feature vectors were used for representation of the biochemical parameters of HVR1.

For the biochemical parameter-based identification of chronically and recently infected hosts, the CFS-based binary feature vector representation of the intra-host HCV population was used as input data to train a stochastic gradient descent (SGD) classifier^26,8^ as implemented on Scikit-learn version 0.20.0). The SGD classifier implements regularized linear models with stochastic gradient descent (SGD) learning and is a very efficient approach, with linear training cost, which can easily be scaled to big data problems (more than 10^5^ training samples and 10^5^ descriptors). Selection and tuning of the hyperparameters of the SGD classifier was done using a balanced training set (1,965 CFS-based binary feature vector representations – persistent class– and 1,968 –recent class–), and the GridSearchCV module in Scikit-learn. The parameters of the selected probabilistic logistic regression classifier were: loss function = ‘log’, alpha = ‘0.1’, n_iter = 100, class_weight = ‘balanced’, random_state = 42. Five-fold cross-validation training of the SGD classifier was done using Scikit’s StratifiedKFold function (parameters: n_splits = 5, shuffle = True, Random_state = 42).

The SGD classifier was used to predict the class probability of each HCV haplotype from its feature vector. Host classification was then performed by computing the expected value of the haplotype predictions, weighted by their frequencies: $\bar{x}=\sum_{i=1}^{n} \left( F_{i}\times P_{i} \right)\div\sum_{i=1}^{n} F_{i}$, where *n* is the number of HCV haplotypes obtained from a host, *F*, the haplotype’s frequency and *P*, the predicted class probability.

^S6^Friedel M, Nikolajewa S, Sühnel J, Wilhelm T. DiProDB: a database for dinucleotide proper- ties. Nucleic acids research. 2008;37(suppl_1):D37–D40.

^S7^Liu B, Liu F, Wang X, Chen J, Fang L, Chou KC. Pse-in-One: a web server for generating various modes of pseudo components of DNA, RNA, and protein sequences. Nucleic acids research. 2015;43(W1):W65–W71.

^S8^Tsuruoka, Y., Tsujii, J.I. and Ananiadou, S., Stochastic gradient descent training for l1-regularized log-linear models with cumulative penalty. In *Proceedings of the Joint Conference of the 47th Annual Meeting of the ACL and the 4th International Joint Conference on Natural Language Processing of the AFNLP.* 2009(1), 477-485.

^S9^Kurgan LA, Cios KJ. CAIM discretization algorithm. IEEE transactions on Knowledge and Data Engineering. 2004;16(2):145–153.

^S10^Wang Y, Tetko IV, Hall MA, Frank E, Facius A, Mayer KFX, et al. Gene selection from microarray data for cancer classification--a machine learning approach. Comput Biol Chem. 2005 Feb;29(1):37–46.
